# A simple bypass assay for DNA polymerases shows hypermutating variants associated with cancer show mechanistic differences in vitro

**DOI:** 10.1101/2022.01.10.475213

**Authors:** Gilles Crevel, Stephen Kearsey, Sue Cotterill

## Abstract

Errors made by DNA polymerases contribute to both natural variation and, in extreme cases, to genome instability and its associated diseases. Recently the importance of polymerase misincorporation in disease has been highlighted by the identification of cancer-associated polymerase variants and the recognition that a subgroup of these variants have a hypermutation phenotype in tumours. We have developed a bypass assay to rapidly determine the tendency of a polymerase to misincorporate in vitro. We have used the assay to compare misincorporation by wild-type, exonuclease defective and two hypermutating DNA polymerase e variants, P286R and V411L. The assay clearly distinguished between the misincorporation rates of wild type, exonuclease dead and P286R polymerases. However, the V411L polymerase showed different misincorporation characteristics to P286R, suggesting that these variants cause hypermutation by different mechanisms. Using this assay misincorporation opposite a templated C nucleotide was consistently higher than for other nucleotides, and this caused predominantly C to T transitions. This is consistent with the observation that C to T transitions are commonly seen in POLE mutant tumours.

## INTRODUCTION

Accurate synthesis by the replicative DNA polymerases is vital for the maintenance of genomic stability. 98-99% of this synthesis is carried out by DNA polymerases δ and e which have a high degree of accuracy (1) (∼1 in 10^-4^ -10^-6^ from in vitro measurements), the rest being carried out by DNA polymerase a (2), which is slightly less accurate (∼1 in 10^-3^ -10^-4^). The main reason for the increased accuracy of DNA polymerases δ and ε over polymerase α is their possession of a 3’-5’ exonuclease activity, which is able to proofread and remove nucleotides which have been incorrectly incorporated. The exonuclease active site is in the largest subunit for both pol δ and ε (3) (4) (5) (6) (7). This subunit also contains a separate polymerase active site, and the nascent strand terminus moves ∼40Å from the polymerase to the exonuclease site for proofreading. Loss of proofreading ability leads to a higher rate of synthesis errors in vitro and also in vivo in both S. cerevisiae and mice. (1) (8) (9) (10)

Recently mutations have been identified in the pol/exo subunit of both DNA polymerase δ and ε that appear to be causative for the generation of tumours (reviewed in (11) (12) (13). These were originally seen in colon and endometrial cancer, but more recently have also been reported in other cancers. Many of the mutations studied so far have been in the exonuclease domain of the pol/exo subunit, and so their effects could be explained by the lack of exonuclease activity raising the misincorporation rate higher than can be handled by downstream mechanisms (particularly MMR -mismatch repair). However, for some mutations the misincorporation observed in vivo is higher than that observed for an exonuclease dead mutant. This suggests that factors other than the lack of exonuclease activity must be responsible for the high mutation rates observed by these polymerases.

Many factors could be responsible for the observed hypermutation. These could be integral to the polymerase itself (eg changes in rate or processivity of synthesis), or could be due to factors only occurring in vivo (eg change in interaction with other proteins or due to handover to polymerases with lower fidelity). When studying the mechanisms of mutant polymerases it is useful to know which of these possibilities is responsible for the high error rate, as this will give insight into how high replication fidelity is normally achieved.

P286R and V411L are the two Polymerase ε (POLE) variants that are the most frequent somatic variants associated with hypermutated cancer cells (11). P286 is highly conserved in ε polymerases from a wide range of organisms, and also in DNA polymerase δ and phage polymerases (Supplementary figure1A). Yeast cells expressing P286R also show a dramatic hypermutation phenotype. In vitro exonuclease assays show that an N terminal fragment of the human POLE catalytic subunit carrying the P286R mutation shows no exonuclease activity, while similar assays using the S. cerevisiae holoenzyme equivalent to P286R reported reduced but significant activity (14). V411 is also widely conserved in ε polymerases, but not in other polymerases, even those as closely related as δ polymerases (Supplementary figure1B). In vitro exonuclease assays suggest that human V411L retains a low level of activity (15). Curiously, despite the high mutation rates in human cells, the in vivo mutation rate of the yeast V411L variant is similar to that of the wild-type enzyme (16).

Here we present a bypass assay to compare the in vitro misincorporation tendencies of wildtype and mutant polymerase variants, where omission of a specific nucleotide forces the polymerase to misincorporate or generate an indel to synthesise a full-length product. Application of this assay to analyse misincorporation for the P286R and V411L variants of the human POLE holoenzyme are consistent with recent results suggesting that these two variants produce misincorporation via different routes.

## RESULTS

### Wild type DNA polymerase ε has limited ability to bypass a templated A nucleotide when dTTP is not available in vitro

One measure of intrinsic fidelity is to determine how easily a polymerase can bypass a base in the template if the complementary dNTP is not available. This approach has been used previously with POLE in stopped flow measurements (8, 8), and also to determine the dissociation frequency of yeast POLE during the transfer of the primer end between the polymerase and exonuclease sites (17). We therefore designed a template that had only a single A nucleotide present in the sequence, and in which the primer strand was labelled with the infrared dye C800 (selected because it can be analysed by infrared scanning in a quantitative way). The basis of the assay is shown in Fig 1A. In the assay there is a high ratio of primer/template to polymerase, so if the polymerase dissociates at the bypass position it is unlikely to reassociate with the stalled product. In this way the full-length product is likely to reflect misincorporation by the polymerase at the bypass position, followed by continued synthesis in the absence of dissociation.

**Figure 1A.**
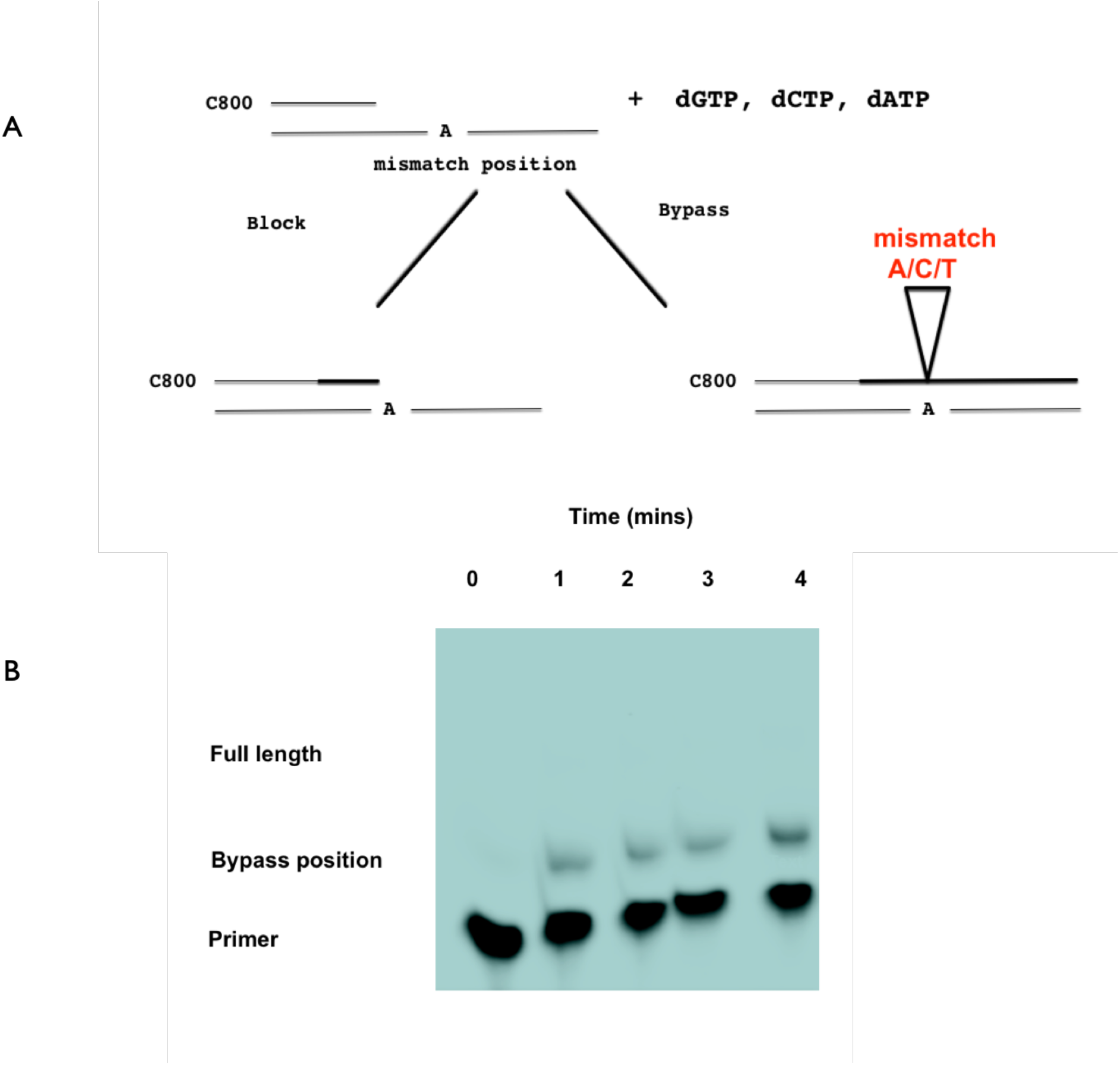
Basis of the Bypass assay. The polymerase is provided with a template that has one of the bases at a single position, (in this case an A – marked as the mismatch position). The enzyme can only proceed to the end of the template if a misincorporation occurs, therefore the ratio of the full length to the blocked template will give a measure of the bypass frequency and the ability of the polymerase to misincorporate. The conditions of the assay are arranged such that there is an excess of substrate compared to the enzyme; if the enzyme dissociates it is more likely to bind to a new template than rebind to the original. The assay therefore represents the ability of an enzyme to misincorporate without dissociation from the template. Figure 1B. Synthesis of the A1 template by the wild type POLE in the presence of only dCTP, dGTP and dATP after 1 2 3 and 4 minutes of reaction. The positions of the full length, bypass position and unreplicated primer are shown.

Synthesis carried out on this template by the wild-type enzyme (purified as described in Materials and Methods) with a full complement of dNTPs gave complete synthesis of the template with increasing yield with time (Supplementary figure 2A/B), whereas synthesis carried out in the absence of dTTP showed increasing synthesis with time but most synthesis was stalled at the position of the single A base (Fig1B).

### The exonuclease dead and P286R variants bypass a templated A with higher frequencies than the wildtype enzyme

This encouraged us to use this assay to compare the in vitro bypass rate of the wild-type enzyme with the exonuclease dead and P286R mutants.

We therefore introduced mutations into the large subunit of POLE to generate exonuclease dead and P286R variants. An analysis of the exonuclease activity of the wild-type, and the exonuclease dead and P286R variants (supplementary figure 3A/B)) showed that neither the exonuclease dead or P286R mutants showed exonuclease activity in vitro. This is consistent with previous observations from in vitro studies using an N terminal fragment of the human enzyme (11). This suggests that previously observed differences in exonuclease activity between the yeast and human P286R variants was not caused by the use of a truncated form of the human enzyme and may indicate that the budding yeast enzyme is affected differently by the mutation.

An example gel for the bypass analysis with these 3 variants is shown in Figure 2A. The exonuclease dead enzyme showed a higher bypass rate than the wildtype, and P286R showed a higher rate still. Quantitation of the results for multiple experiments using template A1 are shown in Figure 2B

**Figure 2A.**
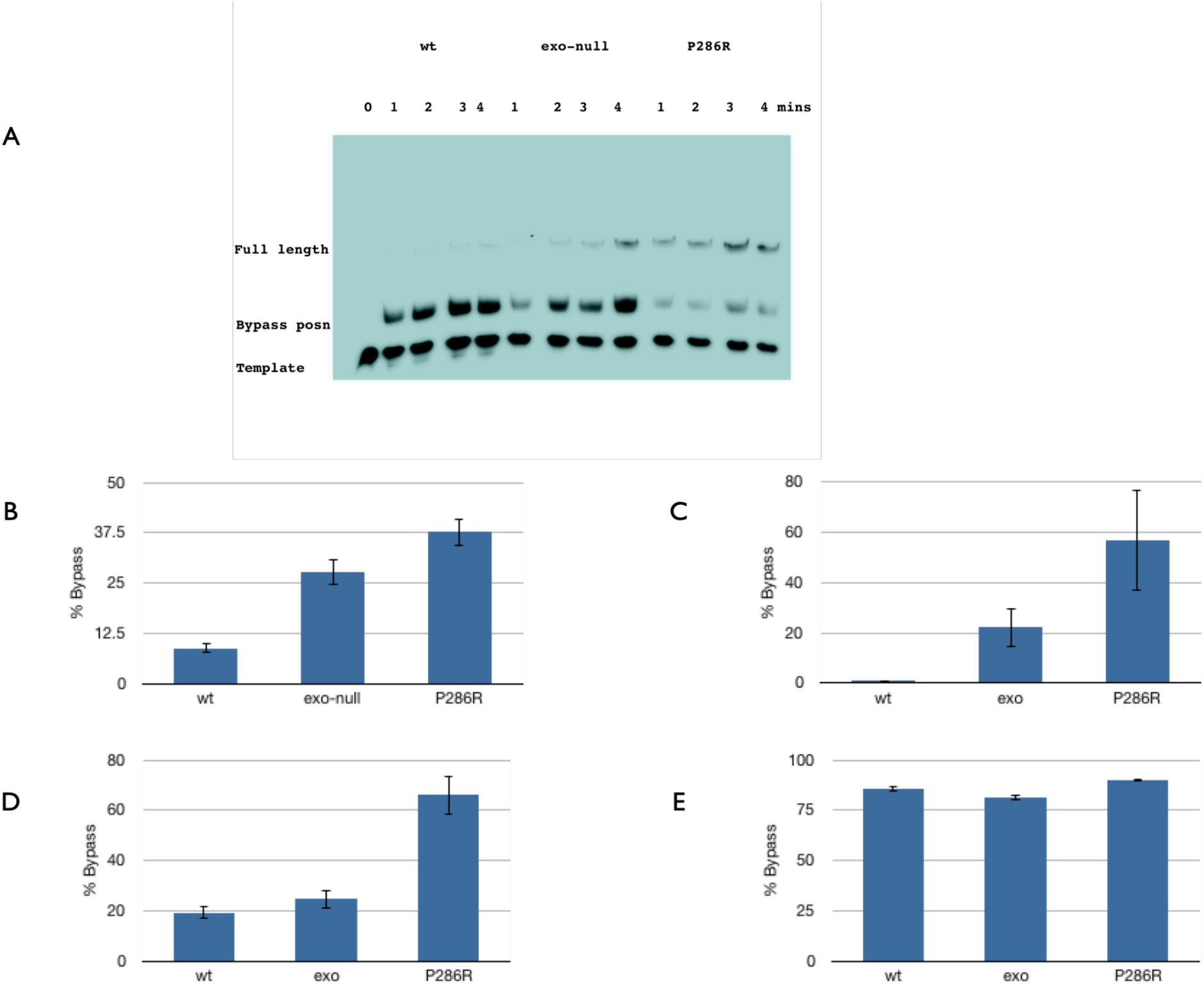
Synthesis of A1 template by wild type, exonuclease dead and P286R variants in the presence of only dCTP, dGTP and dATP after 1-4 minutes of synthesis. 0 is the unreplicated template The positions of the full length, bypass position and unreplicated primer are shown. Figure 2B – Percentage bypass rates on the A1 template for the wt, exonuclease dead and P286R variants in the presence of only dCTP, dGTP and dATP. The error bars represent the standard deviation in the data. Figure 2C Percentage bypass rates for the G template for the wt, exonuclease dead and P286R variants in the presence of only dATP, dGTP and dTTP. The error bars represent the standard deviation in the data. Figure 2D Percentage bypass rates for the T template for the wt, exonuclease dead and P286R variants in the presence of only dCTP, cGTP and dTTP. The error bars represent the standard deviation in the data. Figure 2E Percentage bypass rates for the C1 template for the wt, exonuclease dead and P286R variants in the presence of only dCTP, cATP and dTTP. The error bars represent the standard deviation in the data. For all the above experiments the data presented is derived from 3 independent experiments each with 4 time points.

Comparison of the bypass rate at multiple timepoints (supplementary figure 4) showed that the percentage bypass did not increase with time, provided that a significant fraction of the template remained unreplicated. This implies that the stalled product, once generated, is not eventually extended to full length, suggesting that the full-length product is likely to be due to a misincorporation event which is extended by the same polymerase without dissociation. Therefore, in all subsequent experiments bypass rates presented are an average of the bypass at each time point for each enzyme.

These results suggest that this assay is sufficiently sensitive to detect differences in the bypass rates between polymerases with different fidelities. It further suggests that the hypermutation phenotype of the P286R mutant is at least in part amenable to in vitro analysis, as it has higher bypass rate in the in vitro analysis than the exonuclease dead mutant.

### Differences in bypass efficiency for POLE mutants are also observed for bases other than A

Similar experiments were then carried out to determine whether differential bypass by wild type, exonuclease dead and P286R polymerases could also be observed for the 3 other nucleotides. For both G (Figure 2C) and T (Figure 2D) templates a similar trend was observed although the absolute bypass rates varied for different nucleotides. For the C1 template however, a very high bypass rate was observed even with the wild-type enzyme (fig 2E). This shows that differences in misincorporation between wild-type exonuclease dead and P286R enzymes can be seen for dGTP and dTTP using this assay.

### Bypass rates of ‘single C’ templates are influenced by the context of the base and the concentration of dNTPs in the reaction

The very high rate of bypass on the C1 template was surprising, especially in the case of the wild-type enzyme, but was entirely reproducible using different independent batches of nucleotides and several independently synthesised templates carrying the same sequence. Comparison of the results observed using the A1, G and T templates suggested that although the relative abilities of the different POLE enzymes to misincorporate opposite the single base were conserved, there were differences in absolute bypass rates between different bases. This could be intrinsic to individual bases, however the absolute bypass rate could also be influenced by the context of the bypassed base.

We therefore determined how the ‘single C’ bypass rate was affected by the context in which the C is embedded. We synthesised a template (C2) in which the immediate environment of the single C residue was altered 4 bp upstream and downstream of the C. This change did not affect the base composition of the template but made the 2 bases either side of the C more AT rich (GGAAGT<u>C</u>GTATTA to GGGTAA<u>C</u>TTAGTA). For this template the bypass rate was slightly reduced, (Figure 3A), but still remained much higher than for the other nucleotides. We also made an additional C template (C3) where the downstream sequence was as the original template, but the upstream template was more substantially altered. In addition the single C was moved further away from the primer terminus by the insertion of 8 nucleotides (TTGAAGTAGTGAGATGGAAGT<u>C</u> to TTGAAGTAGTGAGATGGAAGTGTAGATTA<u>C</u>). This template also produced bypass levels for all 3 enzymes that were slightly lower than those for the other C templates, (Figure 3A), but again bypass levels, particularly for the wild-type enzyme were higher than for other bases.

**Figure 3A.**
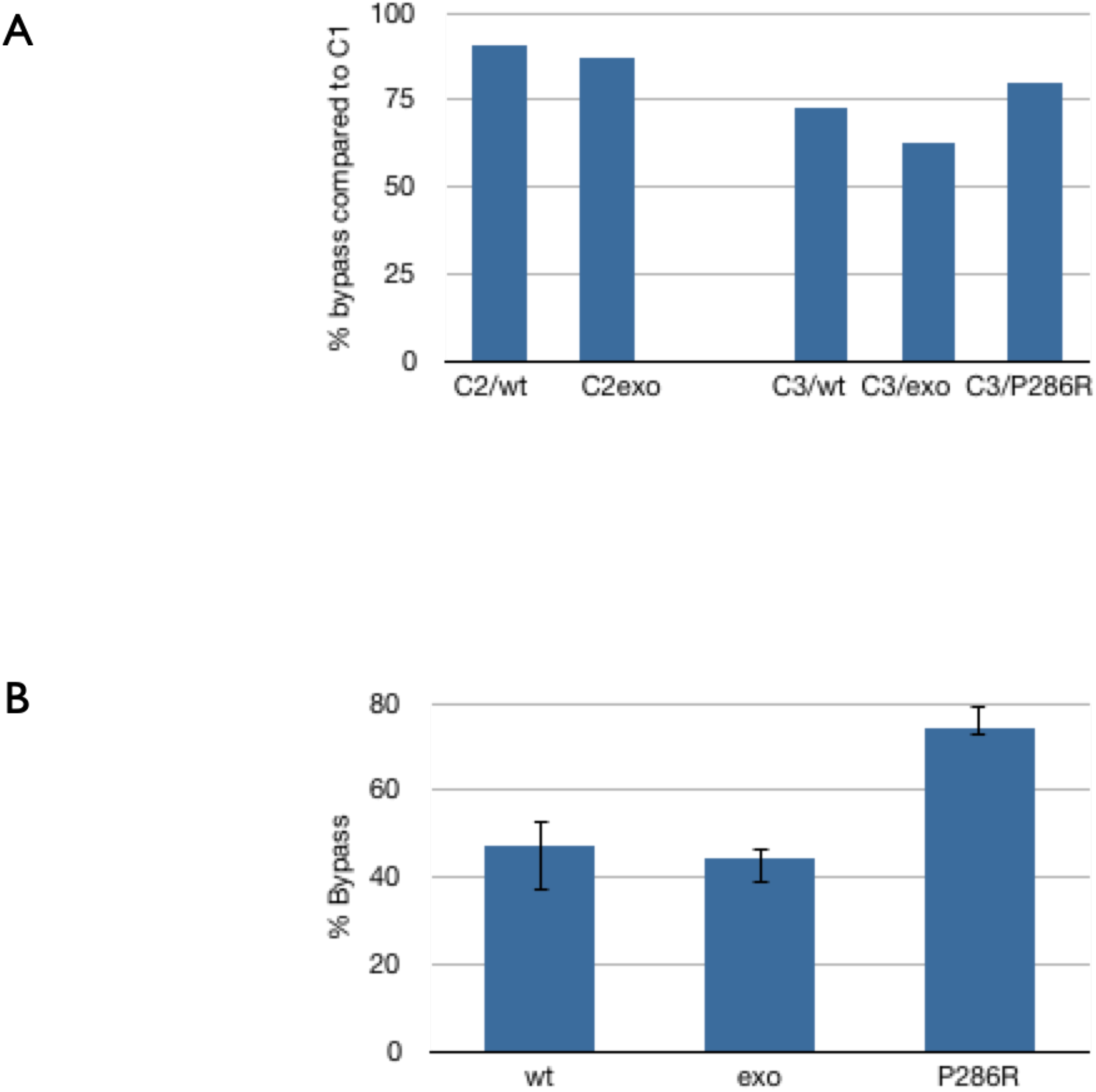
Comparison of bypass rates for templates C1, C2 and C3 at 200uM dATP, dCTP and dTTP. The Y axis shows the ratio of the bypass rate of C2 and C3 compared to C1. The X axis shows the template and the variant used for the assay: wt – wild-type, exo-exonuclease dead and P286R – P286R. Figure 3B Percentage bypass rates on the C3 template for the wt, exonuclease dead and P286R variants in the presence of dCTP, cATP and dTTP at 80uM. These results are taken from 3 independent experiments each with 4 time points. The error bars represent the standard deviation in the data.

Another factor that could influence the amount of bypass is the concentration of the dNTPs present in the reaction. The presence of an increased concentration of dNTPs encourages misincorporations in vivo in a variety of organisms and this is likely to be at least partly due to direct effects on the polymerase (18).Figure 3B shows that reducing the concentration of dNTPs in the reaction by 60% (to 80mM) reduced bypass levels quite significantly for the C3 template. However, bypass levels still remained higher for C3 at low nucleotide concentrations than for the A1, G and T templates at higher dNTP concentrations. Again, this was particularly noticeable for the wild-type enzyme.

One curious feature that we observed consistently for bypass on all C templates at all dNTP concentrations was that the exonuclease dead enzyme had a similar bypass rate to the wild-type enzyme although for the C3 template the P286R read through is clearly higher. These results suggest that it is possible to optimise the bypass assay for a ‘C’ template so that differences in bypass can be observed between a wild-type enzyme and a hypermutating enzyme, but in its present format, bypass of a single C may not be able to distinguish a wildtype enzyme from one that has only lost exonuclease activity.

### Bypass on the C3 template predominantly causes a G to A (C to T) misincorporation

It still remained a formal possibility that some contamination in either the template or the dNTP mix was responsible for the high bypass rate with the C templates. Ganai et al (17) showed that for purified yeast POLE holoenzyme, very low levels of dGTP (0.34-0.68 uM) were enough to support some synthesis. In order to determine whether this was the case we developed a protocol to sequence the DNA that had been synthesised by the POLE variant enzymes. This protocol allows us to use very small amounts of synthesised product and so would be applicable to reactions where the bypass is less efficient. The scheme uses cycle sequencing to determine the base that had been incorporated at the single C position. The scheme is shown in fig 4A and incorporates steps to remove non reacted or partially extended primers, and also as much as possible of the original template, all of which interfere with the cycle sequencing reaction if they remain in the solution at a significant level.

**Figure 4.**
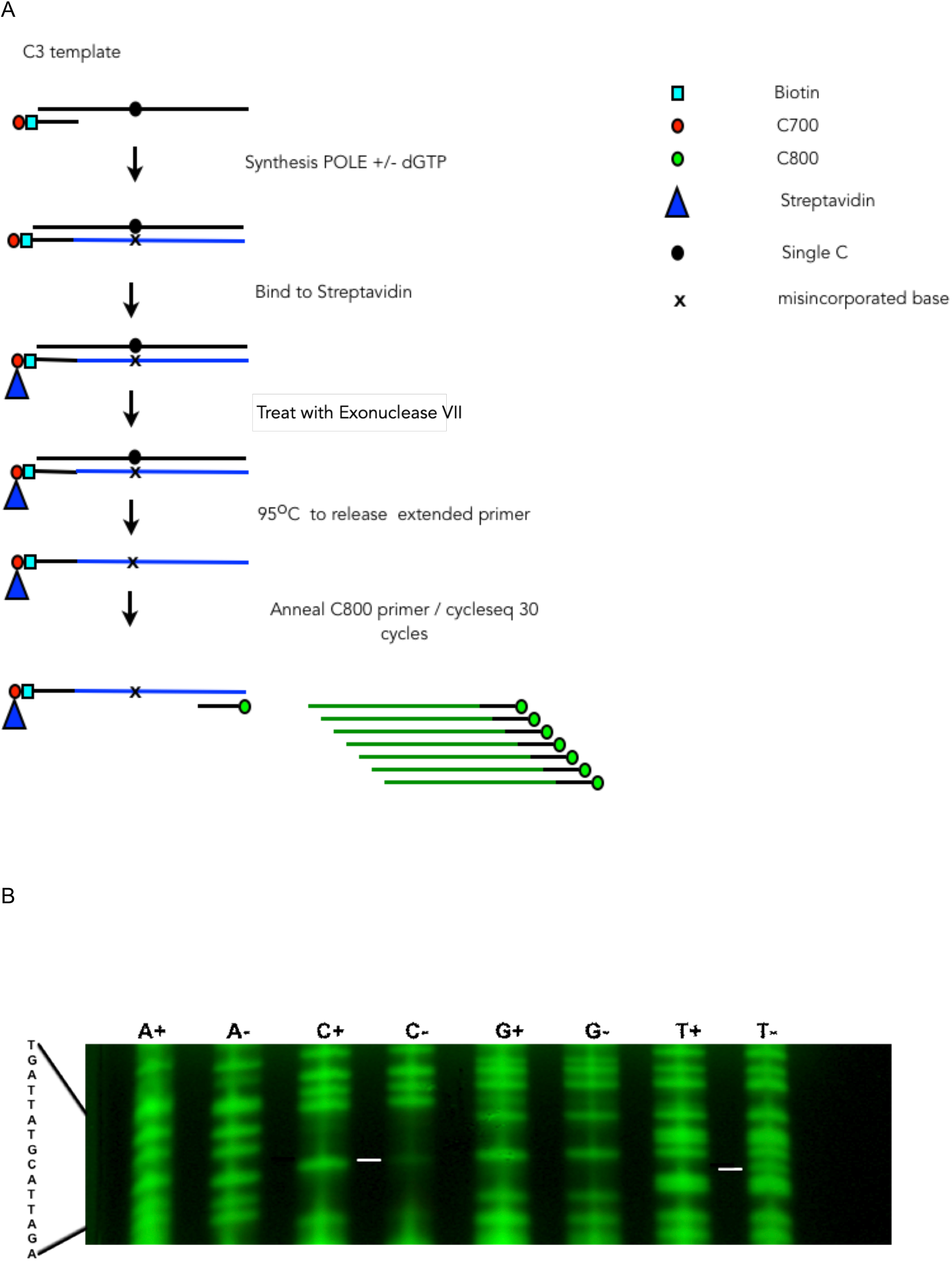
A shows the scheme for sequencing of the products of POLE synthesis. The presence of the C700 label in the biotin labelled primer allows verification of the extent of bypass by POLE in the initial reaction. The exonuclease VII step is designed to remove unannealed template and unextended primer. The heating step is designed to allow removal of the template strand. Figure 4B – shows the results from the cycle sequencing following the bypass assay using the C3 template and the P286R variant. The equivalent nucleotide lanes are run next to each other to make it easier to see changes. A C G T show the chain terminator used for sequencing, + is the analysis of the bypass reaction containing all 4 nucleotides,, - is the analysis of the bypass reaction missing dGTP from the extension mix. The expected sequence is shown at the left hand side of the gel. The lines in the gel show the position of a visible change between the reaction containing all 4 nucleotides and the one missing a G in the extension mix., comprising a reduction in the C- lane and a new band appearing in the T- lane.

Fig 4B shows the results of a sequencing reaction with the synthesis products of P286R on the C3 template. C3 was used as the template as the single C was far enough away from the sequencing primer to identify the incorporated nucleotide clearly. At the position on the template where the C is incorporated in a reaction containing all 4 dNTPs (+), the bypass reaction (-) shows only trace incorporation of C. This is most likely due to remaining contamination of primer or template from the original reaction. Instead, a new band appears in the T track suggesting that if P286R is unable to incorporate a dG in the template opposite a dC it preferentially incorporates a dA. No new bands are visible in other lanes but we cannot rule out the possibility that bands are present but below the level of detection. Similar results were obtained with both the exonuclease dead and the wild-type enzymes. Skipping (deletion) at the bypass position does not occur as the +/- bands are in alignment on both sides of the bypass position

This confirms that the high bypass rate on the C template cannot be explained only by a contamination of either the template or the dNTP mix. It further suggests that there is some specificity to the dNTP mis-incorporated during the bypass. The apparent ease with which POLE incorporates dA when there is a template dC may account for the frequency of C>T transitions in yeast strains expressing an exonuclease defective POLE in the background of MMR deficiency (19).

### V411L does not show an increased bypass rate compared to the exonuclease dead variant

Another common cancer-related POLE variant that has been identified as a hypermutator is V411L. The location of this mutation is further away from the exonuclease active site than P286R. In addition evidence from studies on human cancers and cancer cell lines carrying different POLE mutations suggests that there may be some differences in the mutational signatures of P286R and V411L in vivo (20) (21) (22). V411L has also been shown to retain some exonuclease activity when the N terminal domain of the large subunit was assayed in vitro (15).

We were interested in determining whether we could detect an increased misincorporation frequency for this variant in vitro. We therefore introduced this mutation into the large subunit of polymerase ε. Consistent with previous results we saw that the holenzyme containing the mutation retained a low level of exonuclease activity (Supplementary Figure 3C). We then analysed the ability of the V411L mutant enzyme to perform bypass synthesis for A1, C1, G and T templates. Although V411L is known to be a hypermutator in human cells, the bypass frequency appeared more similar to that of the exonuclease dead mutation than that of P286R (Fig 5). This suggests that consistent with in vivo observations, the mechanism causing hypermutation in V411L is different to that in P286R.

**Figure 5.**
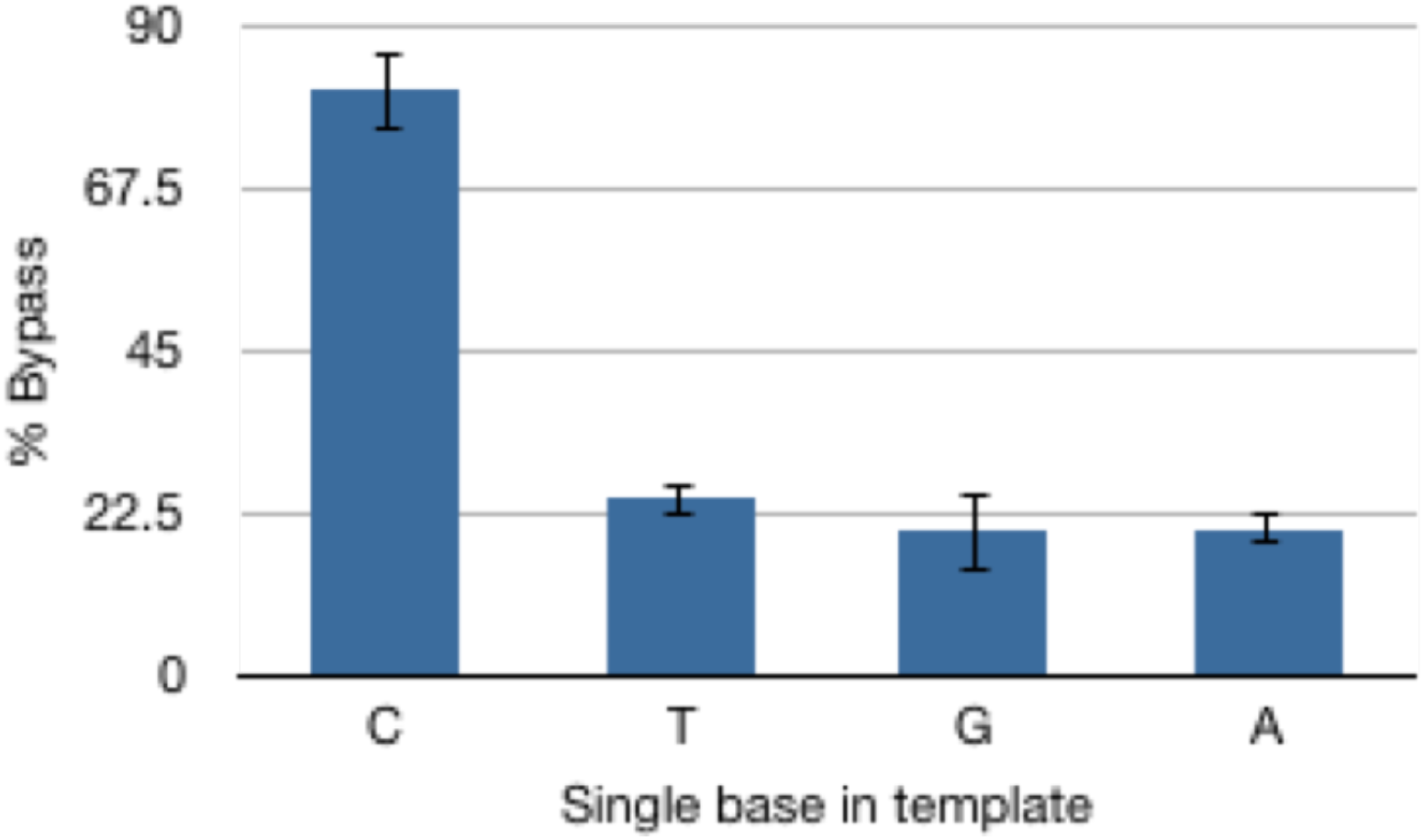
Percentage bypass rates for V411L for the A1, C1, G and T templates. C T G A show the single bases present in each template and in each case the complementary dNTP has been omitted from the reaction. These results are taken from 3 independent experiments each with 4 time points

## DISCUSSION

We have developed a rapid bypass assay to measures the likelihood of polymerase mis-incorporation using templates containing a single A, C, G or T when the required complementary nucleotide is not present in the polymerization reaction. We have used the assay to measure the bypass frequency associated with a 4 subunit wild-type human DNA POLE, and compared this to the bypass observed for a variant of the polymerase that has been mutated to inactivate the exonuclease activity, and 2 other variants that have been altered to contain mutations commonly detected in colon cancer associated polymerases.

Using this assay, it is possible to clearly detect a measurable difference in mis-incorporation ability between purified POLE variants with exonuclease domain mutations. Bypass rates can be determined for a single A G or T in the template. However, the assay does not provide a quantitative measure of in vivo fidelity, as the conditions used for the assay are not conditions that would usually be experienced by the enzyme in a cellular setting. In addition, the precise reaction conditions also affect the amount of bypass that is obtained.

The observed bypass rate in our experiments varies depending on the identity of the templated bases. For instance, the observed bypass ratio of exonuclease dead, P287R and V411L compared to wild type all appear much higher for G than for A, T and C. This could suggest that the wildtype enzyme more accurately incorporates opposite a templated G. However, it is possible that the accuracy observed for the G template could be specific for the particular template and conditions used in the assay. This would be consistent with the data that we obtained using C templates which suggests that the bypass rate might vary depending on the context of the base, and also the concentration of dNTPs that are used in the assay. Similar effects of chromosome context and dNTP concentration have been previously reported (reviewed (23). The concentrations of dNTPs used here, while comparable with the concentrations routinely used for in vitro polymerase assays, are higher than previously measured cellular dNTP levels. These range between ∼5 and ∼40uM depending on the nucleotide (24), but also show differences between different cell types and may also be affected by subcellular location. Direct quantitation of in vivo fidelity rates from the bypass rates obtained with this assay are further complicated as some component of the in vivo measured fidelity could be caused by a corrupted interaction of the polymerase with other cellular components or other factors, such as polymerase switching.

Although not a direct measure of in vivo fidelity, the relative bypass rate measured by this assay does give insight into the likelihood of a polymerase having a mutator phenotype in vivo. Further optimisation of the assay, by altering the template and the nucleotide concentration, should also increase the sensitivity of the assay to allow detection of smaller differences in mis-incorporation frequency and to determine whether the polymerase stalls pre or post misincorporation.

### Distinct behaviour of single-C templates compared to single –A/G/T templates

The bypass assays using the ‘single C’ template showed two features that differed from the other single base templates. The first of these was that all C templates had a much increased level of bypass at the standard dNTP (200uM) concentrations used for the assay compared to A, G and T templates. The C1 template has a TCG context around the C whereas the other 2 templates have ACT (C2) and ACG (C3). TCG is one of the triplets associated with a high frequency of C>T transitions in POLE tumours (21), which could explain why this template had a slightly higher rate of bypass than the other templates. However, the bypass frequency was also high for the other two templates, so this cannot fully explain the high bypass rates observed on ‘single C’ templates.

Reducing the dNTP concentrations to 80uM significantly reduced the bypass rate for the wildtype and exonuclease dead enzymes, although the rate was still higher than that for the other bases at 200mM. This suggests that under the conditions of the assay POLE is more likely to mis-incorporate opposite a C base in the template than opposite any of the other bases. At this point it is not clear why this should be the case. C is the only base that shows substantial methylation in some species, therefore it is tempting to speculate that the relaxed precision may be due to the need to accommodate the methylated variant. In this regard it will be interesting to see whether higher C bypass rates are also found in species which do not show high levels of C methylation.

Sequencing of the POLE synthesised DNA strand from the C3 template showed that there was a very high tendency for an A to be incorporated by the enzyme in place of the G, ie misincorporation results in a C>T transition. A and G are both purines, and so this substitution would cause the minimum disruption to the double stranded structure produced. This substitution is also consistent with the observation that C to T transitions are commonly seen in POLE mutant tumours, particularly (21) in a CG context, in yeast exonuclease-defective POLE strains lacking MMR and in S. pombe expressing the equivalent mutation (in this case P287R) (25). The choice to incorporate an A rather than any other nucleotide must be made at the level of the polymerase catalytic site since the same misincorporation is seen, not only for the wild-type POLE, but also the exonuclease dead and P286R. Neither exonuclease dead or P286R possess exonuclease activity, so it would not be able to change the nucleotide added to the growing chain after incorporation.

The second and more surprising observation was that although the ‘single C’ templates and conditions used here clearly distinguish between a POLE wild-type and P286R mutant they do not allow a definitive distinction between wild-type and exonuclease dead enzymes. For all C templates, the exonuclease dead enzyme showed a similar bypass rate compared to the wild-type enzyme. The reasons for this are not clear, however the wild-type bypass frequency is increased to levels at least as high as that seen with other nucleotides for the exonuclease dead. It is therefore possible that, at least under the specific conditions of this assay, that wild-type POLE may have an impaired capability to remove a misincorporated base opposite a C in the template.

### Hypermutation caused by V411L and P286R may be generated by different mechanisms

We have also used the assay to compare the bypass rate associated with two mutations that are commonly seen in human tumours, P286R and V411L. These variants have been reported to show some differences in mutation signatures in patient samples and cell lines, such that SBS 10B and SBS10A are more predominant in V411L and P286R respectively, and V411L may be more viable if MMR is inactivated (20) (21) (22). In addition, our assays, and those reported by Shinbrot et al (15), show that V411L retains some exonuclease activity, while P286R does not (Supp Fig 3B, C). In contrast to P286R, V411L showed a bypass rate that was similar to the exonuclease dead. This suggests that P286R and V411L generate mutations using different mechanisms. This is not surprising given their different locations. P286R is located at the edge of the exonuclease active site and structural analysis of the yeast enzyme suggested that the amino acid change blocks entry of the nascent DNA terminus to the exonuclease site (26) (14). V411 is quite well separated from the exo site, and lies in the junction region between the exo and polymerase site. It is therefore less likely to block entry to the exonuclease site. It does however lie in a region of the enzyme that could make contacts with the DNA substrate in addition to the active site contacts. Perhaps the position of the V411L substitution makes the polymerase more likely to dissociate from the template as it transfers between the polymerase and exonuclease sites when the wrong nucleotide is incorporated. With the excess of template present in the assay, rebinding would be more likely to occur at a new primer-template than primer-template that is already partly extended and mismatched.

## Conclusion

We have developed an assay that allows us to rapidly measure the intrinsic ability of a purified polymerase to misincorporate in the absence of other accessory factors. This assay should be useful to give an idea of the likely fidelity of mutations in purified polymerase enzymes and will be useful for rapid analysis of cancer-associated mutants to guide routes for further study. Sequence analysis of the misincorporations will also yield insight into the relative contributions to mutation signatures of the properties of the polymerase and downstream repair related activities. Finally, the assay may also be useful as a tool for development of drugs that affect the accuracy of a polymerase, either positively or negatively.

## MATERIAL AND METHODS

### Manufacture of DNA synthesis substrates

Oligonucleotide primers labelled with C700, C800, and where appropriate, also with biotin, were obtained from IDT and non-labelled template oligonucleotides from Sigma. The substrates were made by mixing equal amounts of primer and template oligos (300pM) in 20mM Tris pH8, 10mM KCl, 0.01mM EDTA. Samples were heated to 100°C for 15 mins and then allowed to cool slowly to room temperature. The extent of annealing was assessed using native TBE polyacrylamide gels. In all cases annealing of the labelled primer was estimated to be more than 99%. The oligonucleotides used in the assays are shown in Table 1.

**Table 1.**
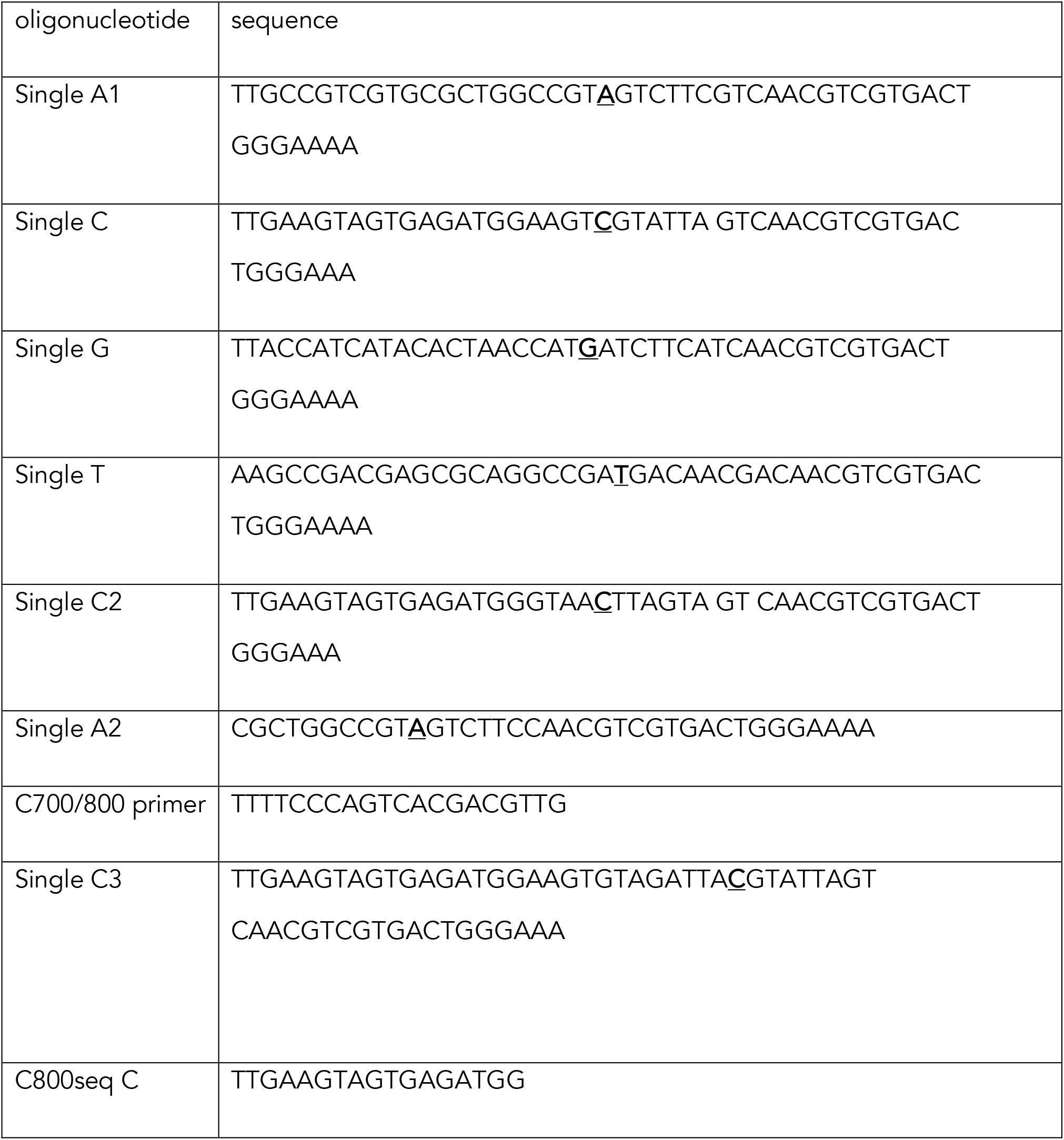
Oligonucleotides used in this analysis:

### Cloning of wild type and mutant DNA polymerases

Clones for the wild type DNA polymerase ε subunits were a kind gift from the Hurwitz lab (17). The original p261 clone was tagged with a flag epitope. Since we were interested in the properties of intact complexes rather than the isolated p261, the tag was removed from the large subunit and a Flag tag added to the p58 subunit. The exonuclease dead, P286R and V411L variants were obtained by carrying out point mutation to these clones using the QuikChange II site directed mutagenesis kit (Agilent). The oligonucleotides used to produce these mutations are shown in Table 2. The changes introduced to make the polymerase devoid of 3’-5’ exonuclease activity (exonuclease dead) were based on work reported from yeast and mice (9) (10).

**Table 2.** Nucleotide changes made to produce the human POLE variants used in this study

| polymerase | Amino acid mutation | Nucleotide mutation |
| --- | --- | --- |
| Wild-type | n/a | n/a |
| exonuclease dead | D275A E277A | a824c<br>a830c |
| P286R | P286R | c857g |
| V411L | V411L | g1231t |

### Purification of wild type and mutant polymerases

The DNA for the wild-type and mutated sequences was introduced into the pFastBac vector (ThermoFisher Scientific) and transfected into DH10Bac E. coli in order to produce bacmids containing polymerase subunits. Selection of positive clones was carried out using blue/white selection. Clones for each of the 4 subunits were used individually to infect sf9 cells to produce baculovirus stocks.

To isolate polymerase complexes Sf9 cells were infected with Bacmids for each of the 4 subunits simultaneously and grown for 72 hours. Cells were harvested and the enzymes were purified as described (27) with a final glycerol gradient step to ensure that only intact holoenzymes were present. The purity of the polymerase enzymes were assessed after the glycerol gradient step by SDS-PAGE and Coomassie gel staining. Protein concentrations were calculated using serial dilutions of Biorad protein markers.

The polymerase activity of each prep was calculated using the A2 substrate in the presence of all 4 nucleotides.

### Bypass assays

Polymerase assays were carried out using templates as described above. 25ul reactions containing enzyme, 12pM DNA template, the appropriate dNTP mix (200mM of each nucleotide), 20mMTris pH 7.5, 10mM magnesium acetate and 150mg/ml BSA were incubated at 37°C. 20fm of wild type polymerase added to reactions to ensure that the template was always in large excess over enzyme. For all assays the amount of each mutant polymerase was adjusted so that it had the same polymerase activity as the wild type enzyme. Samples were taken at the time points indicated. The reaction was stopped by the addition of denaturing gel loading buffer (ThermoFisher) supplemented with EDTA. Samples were heated to100°C for 5 minutes to denature the strands and then analysed on 10% denaturing TBE gels containing 7M urea. The gels were visualised using a Licor Odyssey CLx imaging system and where applicable quantitated using Licor software. In each case the % bypass was calculated as:

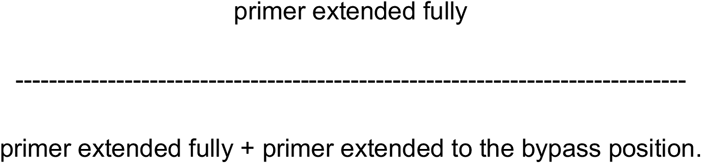

### Exonuclease assays

These were carried out under the same conditions as the polymerase assays except that dNTPs were omitted from the assay, and the samples were run on 15% denaturing gels. The template used was A1 annealed to a version of the standard primer with an added mispaired G on the 3’ end. Where appropriate the exonuclease activity was calculated by quantitation of band intensities using Licor Odyssey software and calculating the total number of bases removed in each case.

### DNA sequencing

DNA sequencing was carried out using the Thermosequenase cycle sequencing kit from Applied Biosystems. Briefly oligonucleotides labelled with C700 and biotin were annealed to the appropriate templates as described. These were used as substrates for a polymerase reaction and the reaction was allowed to proceed for 20 minutes. The reaction was stopped using 25mM EDTA and the products were purified by binding to streptavidin magnetic beads. To reduce background interference from oligonucleotides bound to the column that were unextended or not fully extended at the end of the reaction, the beads were subjected to digestion with exonuclease VII for 60-90 minutes at 37°C. To reduce background interference from the original oligonucleotide templates that remained in the reaction the reactions were subject to 2 rounds of heating to 95°C for 5 mins in TE followed by removal of the supernatant. Following this the polymerase products bound to the beads were sequenced following the instructions from the kit using a primer labelled with C800. The conditions use for the reaction were: anneal: 30s at 52°C, extension: 10s at 70°C and the reaction was carried out for 30 cycles. Amplified products were separated on a 10% denaturing polyacrylamide gels and visualised using a Licor Odyssey scanner. Sequences were read manually.

## SUPPLEMENTARY DATA

Supplementary Data are available at NAR online.

## ACKNOWLEDGEMENT

This work was greatly helped by the gift of expression plasmids from The Hurwitz lab and the authors would particularly like to thank Inger Tappin for help with this.

