## supplementary data for "A simple bypass assay for DNA polymerases shows hypermutating variants associated with cancer show mechanistic differences in vitro"

Supplementary figure 1- Comparison of the sequences of various polymerases in the region of the P286R and V411L mutations

1A – Conservation of the region around P286 in Human polymerase ε . The conserved P is highlighted in red for Human (P286), S.pombe (P287) and S.cerevisiae (P301) ε polymerases and Human (P327) and S.cerevisiae (P332) δ polymerases. The conserved D and E residues which form part of the exonuclease active site are highlighted in blue.

1B - Conservation of the region around V411 in human polymerase ε . The conserved V is highlighted in red for Human (411), S.pombe (411) and S.cerevisiae (426) ε polymerases. This amino acid is not conserved in the δ polymerase family.

Supplementary figure 2 – Normalisation of polymerase concentrations for the assays.

A) Representative gel comparing the synthesis by wild-type, exo-null and P287R POLE variants at 0 1 2 3 and 4 minutes. The positions of the unreplicated primer, the bypass position and the full length are marked. The faint band at the position of the bypass on this gel is due to xylene cyanol which coincidentally runs at the same position and so serves as a good indicator of the position of the template blockage. It can be distinguished from the DNA bands for quantitation purposes due to its different emission wavelength.

B) Reactions similar to the one shown above were carried out for all templates and the results were combined to calculate the relative activity of the enzyme preparations used. In all bypass experiments the amounts of exo-null and P286R enzymes were adjusted so that the enzyme activity present in each reaction was comparable.

Supplementary figure 3. Representative gels showing exonuclease assays for purified Polymerase ε variants .

A) Exonuclease rates for wild-type and exo-null enzymes. C is the unreacted substrate and 1 4 8 and 16 are the time in minutes at which the timepoints were taken.

B) shows the reaction for the P286R variant from the same experiment. In this case the samples have been overloaded and overexposed to check for residual exonuclease activity.

C) shows a comparison of wild-type and V411L variants. C is the unreacted substrate and in this case time points were taken at 1 2 and 4 minutes to allow a more accurate estimation of the level of exonuclease of V411L. Percentage activity for 411 was obtained by repeating the assay 3 times and scanning the gels produced on a Licor odyssey machine.

Supplementary figure 4. Relative bypass rates do not increase with time for the wild-type, exo-null and P286R variants inder the conditions of the assay. The time in minutes is shown on the X axis and the percentage bypass on the Y axis. This shows the results from one assay with the A1 substrate, but similar lack of correlation between bypass rate and time was observed for all assays carried out.

Supplementary Figure 1

## 1A

```
HsPolε    265 ERPDPVVLAFDIETTKLPLKFPDAETDQI 293
SpPolε    266 ERAEPTIMAFDIETTKLPLKFPDSSFDKI 294
T4Pol     102 DRKFVRVANCDIEVTG--DKFPDPMKAEY 128
RB69Pol   104 DHTKIRVANFDIEVTSP-DGFPEPSQAKH 131
HsPolδ    306 RIAPLRVLSFDIECAGRKGIFPEPERDPV 334
SpPolδ    290 KMAPLRIMSFDIECAGRKGVFPDPSIDPV 318
                        ***       **
```

## 1B

```
HsPolε    389 FQKDSQGEYKAPQCIHMDCLRWVKRDSYL 417
SpPolε    390 FFRDAEDEYKSSYCSHMDAFRWVKRDSYL 418
ScPolε    404 FAPDAEGEYKSSYCSHMDCFRWVKRDSYL 432
```

Supplementary Figure 2

A

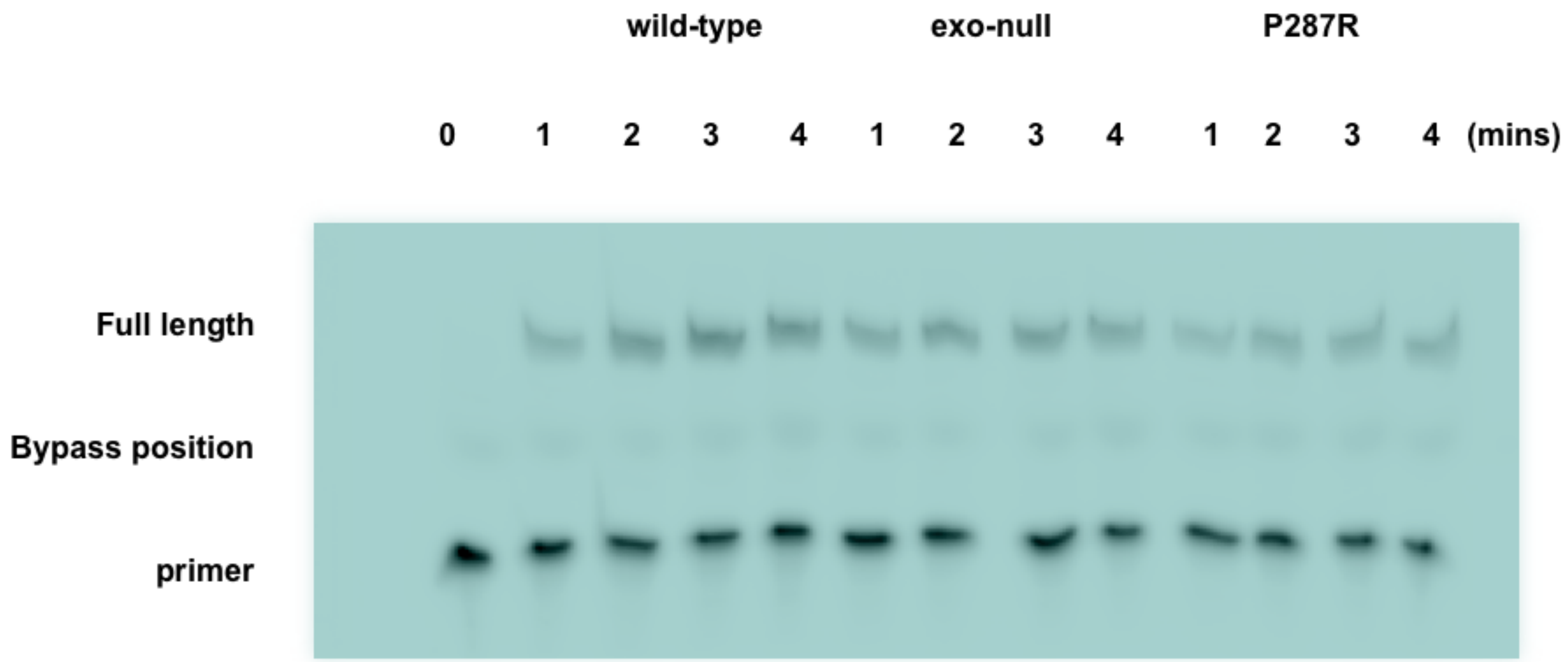


B

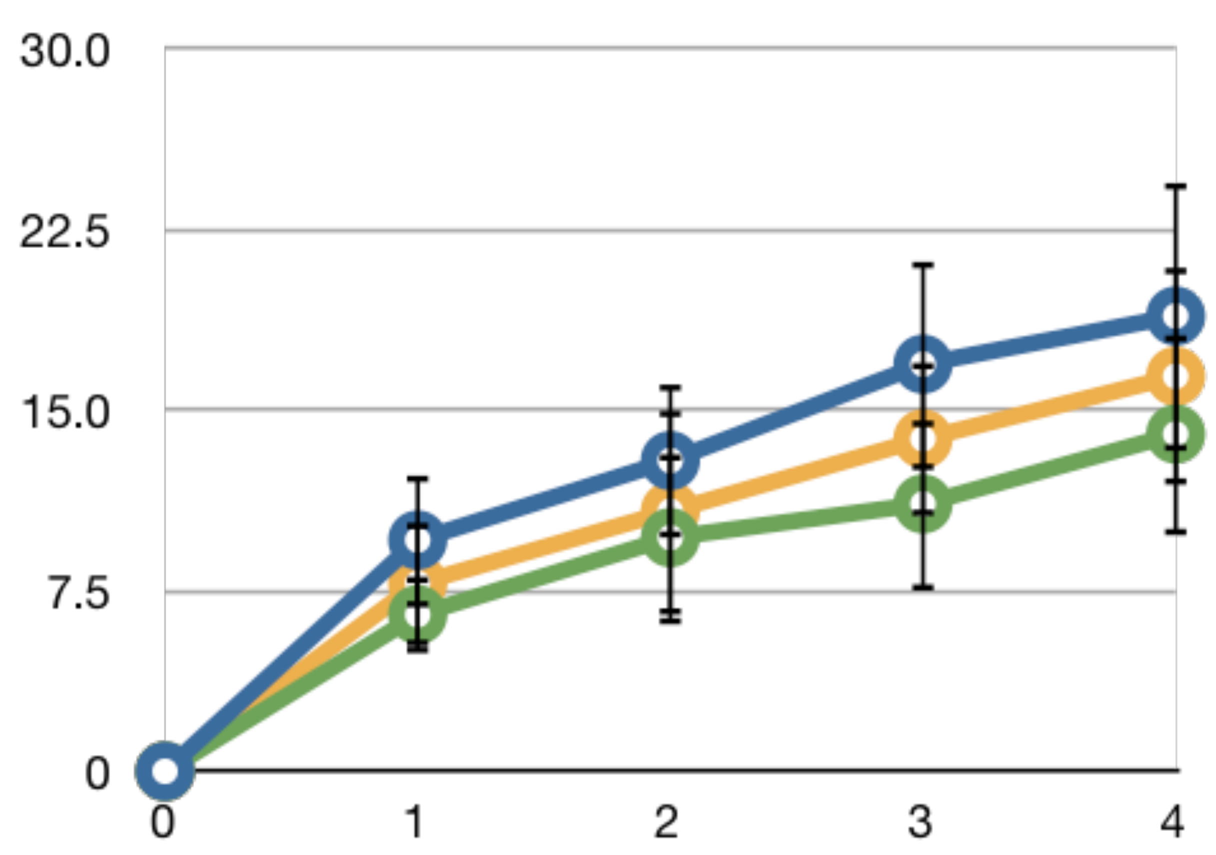

Supplementary Figure 3

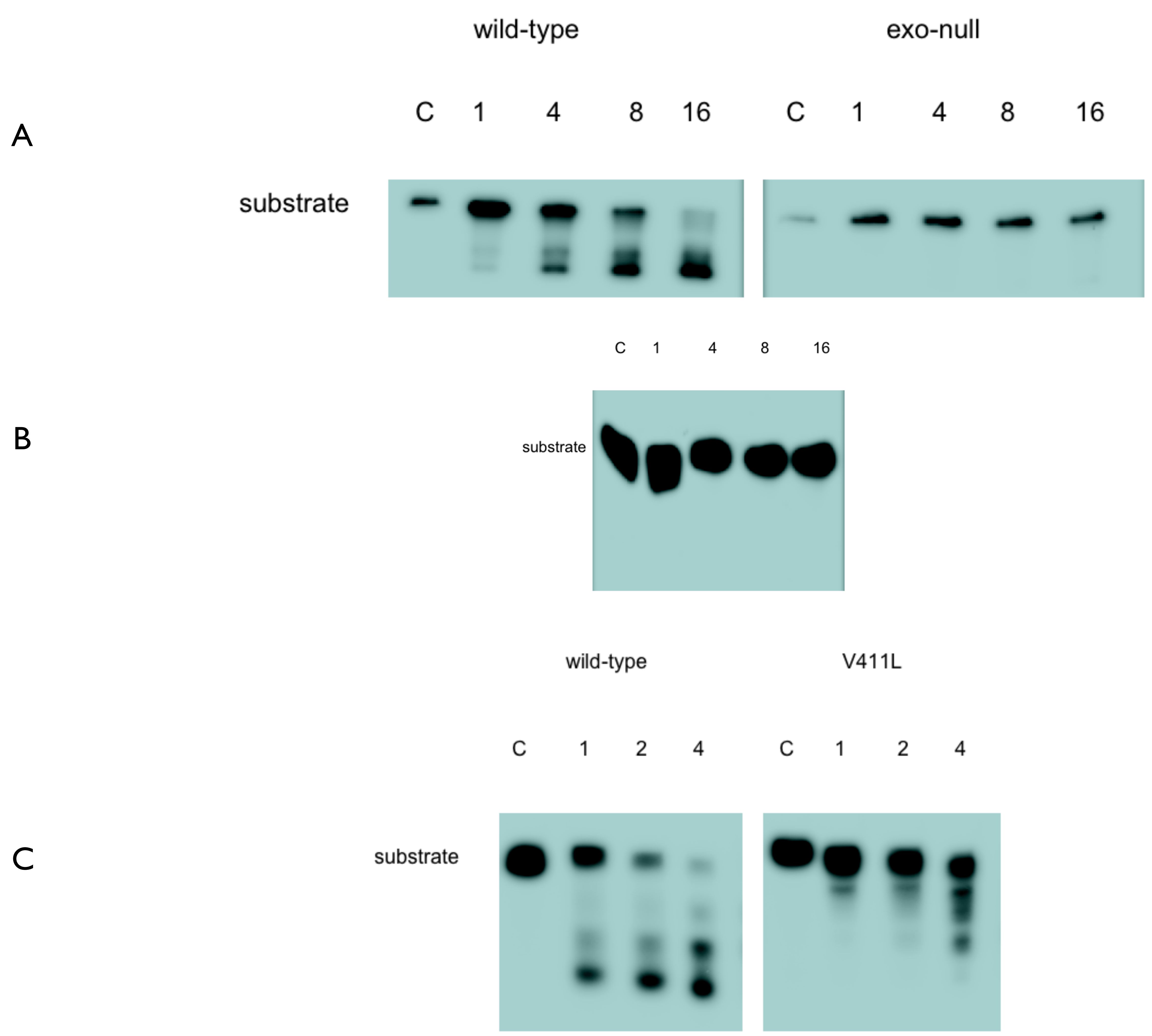

Supplementary Figure 4

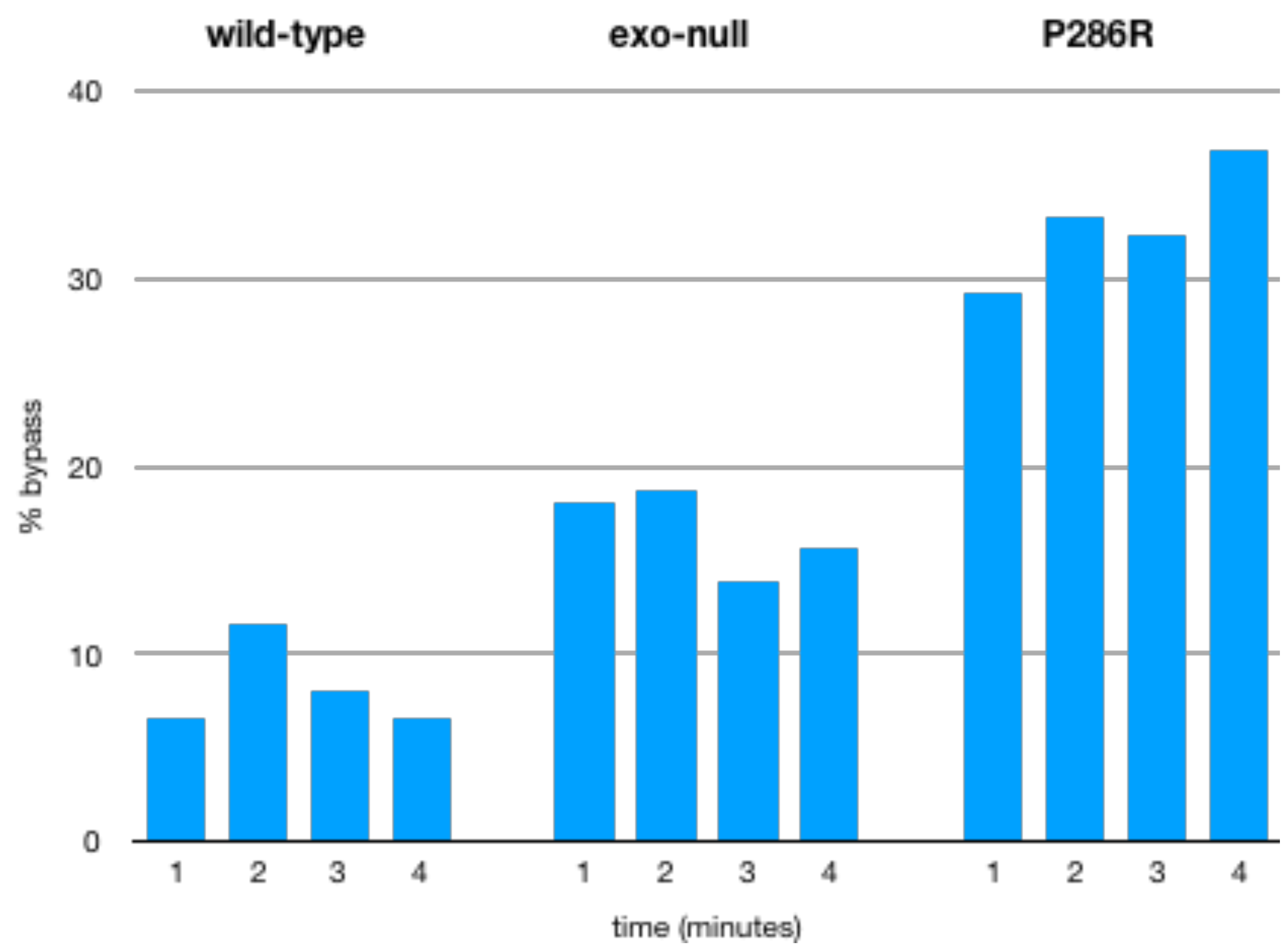
